# PILS6 is a temperature-sensitive regulator of nuclear auxin input and organ growth in *Arabidopsis thaliana*

**DOI:** 10.1101/250001

**Authors:** Elena Feraru, Mugurel I. Feraru, Elke Barbez, Lin Sun, Angelika Gaidora, Jürgen Kleine-Vehn

**Affiliations:** Department of Applied Genetics and Cell Biology, University of Natural Resources and Life Sciences (BOKU), Muthgasse 18, 1190 Vienna, Austria

## Abstract

Global warming is threatening plant productivity, because plant growth is highly sensitive to elevated temperatures. High temperature (HT) triggers the auxin biosynthesis-dependent growth in aerial tissues. On the other hand, the contribution of auxin to HT-induced root growth is currently under debate. Here we show that the putative intracellular auxin carrier PIN-LIKES 6 (PILS6) is a negative regulator of organ growth and that its abundance is highly sensitive to HT. PILS6 localises to the endoplasmic reticulum (ER) and limits the nuclear availability of auxin, consequently reducing the auxin signalling output. HT represses the transcription and protein abundance of PILS6 specifically in the root tip, which impacts on PILS6-dependent root organ growth rates. Accordingly, we hypothesize that PILS6 is part of a novel mechanism, linking HT to auxin responses in roots.

## Introduction

One of the major negative consequences of human progress on the environment is climate change. Climate change, including decreasing cold temperature extremes and increasing warm temperature extremes (IPCC, 2014), is among the biggest threats the world is facing today. The last 30 years were likely the warmest of the last millennium in the Northers Hemisphere and, unfortunately, the global mean surface temperature is predicted to further rise over the 21^st^ century by 1-4°C (IPCC, 2014). The increase in the global surface temperature will subsequently affect soil temperature (IPCC, 2014) and, thus, global warming is becoming a big threat for animals and plants.

An increase in ambient temperature has dramatic consequences on plant development, including crop productivity. *Arabidopsis thaliana* is a suitable model system to investigate adaptation to temperature. The optimal growth temperature for *Arabidopsis thaliana* is 22-23°C (Rivero et al., 2014; ABRC 2013), and is in the range of so called warm ambient temperature (22-27°C; Liu et al., 2015). Temperatures above 27°C are classified by Liu and co-authors (2015) as either high temperature (HT; 27-30°C) or extremely HT (37-42°C), which are levels that strongly affect various aspects of plant development.

Phytohormones are important regulators to integrate external signals into the growth program, allowing for adaptive plant growth and development. The endogenous auxin indole-3-acetic acid (IAA) is a major plant growth regulator (Davies, 2010) and is also fundamentally important for responses to deviation in ambient temperature (Shibasaki and Rahman, 2013; Ahammed et al., 2016).

In aerial organs, such as hypocotyls and petioles, phytochrome B (phyB) functions as a thermoreceptor (Jung et al., 2016; Legris et al., 2016). HT induces the inactivation of phyB, which de-represses the bHLH transcription factor phytochrome interacting factor 4 (PIF4), being crucial for aerial tissues to respond to HT (Jung et al., 2016; Legris et al., 2016). Ultimately, HT-induced PIF4 does elevate auxin biosynthetic genes, which will consequently induce growth in aerial tissues (Gray et al., 1998; Koini et al., 2009; Franklin et al., 2011; Sun et al., 2012).

Compared to the shoot, it remains mechanistically puzzling how elevated temperature impacts upon root growth and development. An increase in temperature (26^o^C-29^o^C) also stimulates primary root growth in *Arabidopsis* seedlings (Hanzawa et al., 2013; Wang et al., 2016; Fei et al., 2017; Martins et al., 2017). However, the underlying hormone-based mechanism is currently under debate. While several studies suggest that HT also affects root growth in an auxin-dependent manner (Hanzawa et al., 2013; Wang et al., 2016; Fei et al., 2017), a recent study shows that brassinosteroid, but not auxin signalling, regulates ambient temperature adaptation in roots (Martins et al., 2017). A central argument in the latter study is that, besides their prominent roles in shoots, PIF4 and its downstream auxin biosynthetic genes do not link temperature sensing with growth responses in roots (Martins et al., 2017).

The PIN-LIKES (PILS) proteins are putative auxin carriers at the endoplasmic reticulum (ER), which limit auxin signaling, possibly by sequestering auxin in the ER (Barbez et al., 2012; Barbez et al., 2013). Notably, the importance of PILS2, 3, and 5 for light-induced growth in apical hook development was recently shown (Beziat et al., 2017), proposing that PILS proteins integrate environmental signals to induce auxin signalling minima. Here we show that PILS6 is a temperature-sensitive regulator of nuclear availability of auxin, contributing to organ growth.

## Results and discussion

Here we specifically characterised PILS6, because it is evolutionarily most distantly related to the so far characterized PILS2, 3, and 5 proteins (Feraru et al., 2012). To assess the importance of PILS6 for seedling development, we isolated the *pils6-1* full knock-out and hypomorphic *pils6-2* mutants in *Arabidopsis thaliana* (Alonso et al., 2003; Supplementary Figures 1A, 1B). Under standard growth conditions at 21°C, the mutation in *pils6* induced overall increased organ growth, displaying bigger rosette as well as cotyledon area, and longer roots (Figures 1A-1E). To assess how constitutive PILS6 expression affects plant growth, we expressed PILS6-GFP under the control of 35S promoter (*PILS6*^*OE*^) and generated several independent, stable transgenic lines (Supplementary Figure 2A). Conversely to *pils6* mutants, *PILS6*^*OE*^ led to smaller rosettes as well as cotyledons, and inhibited primary root growth (Figures 1A-1E). The impact of PILS6 on organ size is likely related to cell size regulation, because the *pils6* mutants and *PILS6*^*OE*^ displayed larger and smaller pavement cells (PC) as well as stomatal cells, respectively (Figures 1F-1H; Supplementary Figures 1C-1E). Altogether, this data indicates that PILS6 is a negative regulator of cell size and organ growth in *Arabidopsis*.

**Figure 1.**
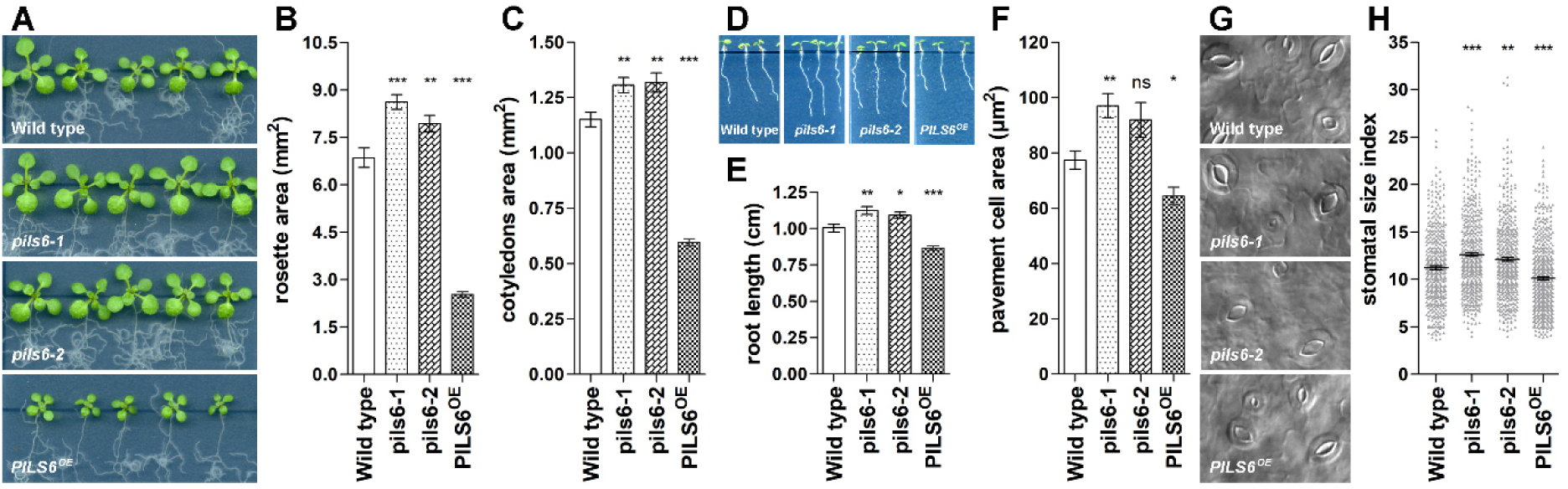
PILS6 is a negative regulator of cell size and organ growth. A-E PILS6 affects organ growth. Scanned images (A, D) or measurements (B, C, E) show that rosette area (A, B), cotyledon area (C) and total root length (D, E) are affected in the *pils6* mutants and *PILS6*^*OE*^ when compared to the wild type control. F-H PILS6 affects cell size. Microscope images (G) and measurements (F, H) show that PC area (F) and stomatal size (G, H) are affected in the *pils6* mutants and *PILS6*^*OE*^ when compared to the wild type control. n= 28 rosettes (B), 50-56 cotyledons (C), 49-55 roots (E), 62-119 PC from 6 cotyledons per each genotype (F), and 850-1092 stomata from 14 cotyledons per each genotype (H); ns: not significant; *: p< 0.05, one-way ANOVA.

Next, we addressed the subcellular localisation of PILS6-GFP in roots. Similar to functional PILS3-GFP (Beziat et al., 2017) or PILS5-GFP (Barbez et al., 2012), transgenic PILS6*-*GFP resided in perinuclear and reticular membranes, indicating localisation at the ER (Figure 2A; Supplementary Figure 2A). Compared to lines overexpressing PILS6-GFP, we obtained a similar, albeit much weaker, PILS6-GFP localisation when expressed under its endogenous promoter (Figure 2B; Supplementary Figures 2B, 2C). Notably, *pPILS6:PILS6-GFP* complemented the *pils6-1* mutant phenotype (Supplementary Figure 2D), indicating that PILS6-GFP protein fusion is functional. Accordingly, our data suggests that PILS6 impacts at the ER on cell and organ size.

**Figure 2.**
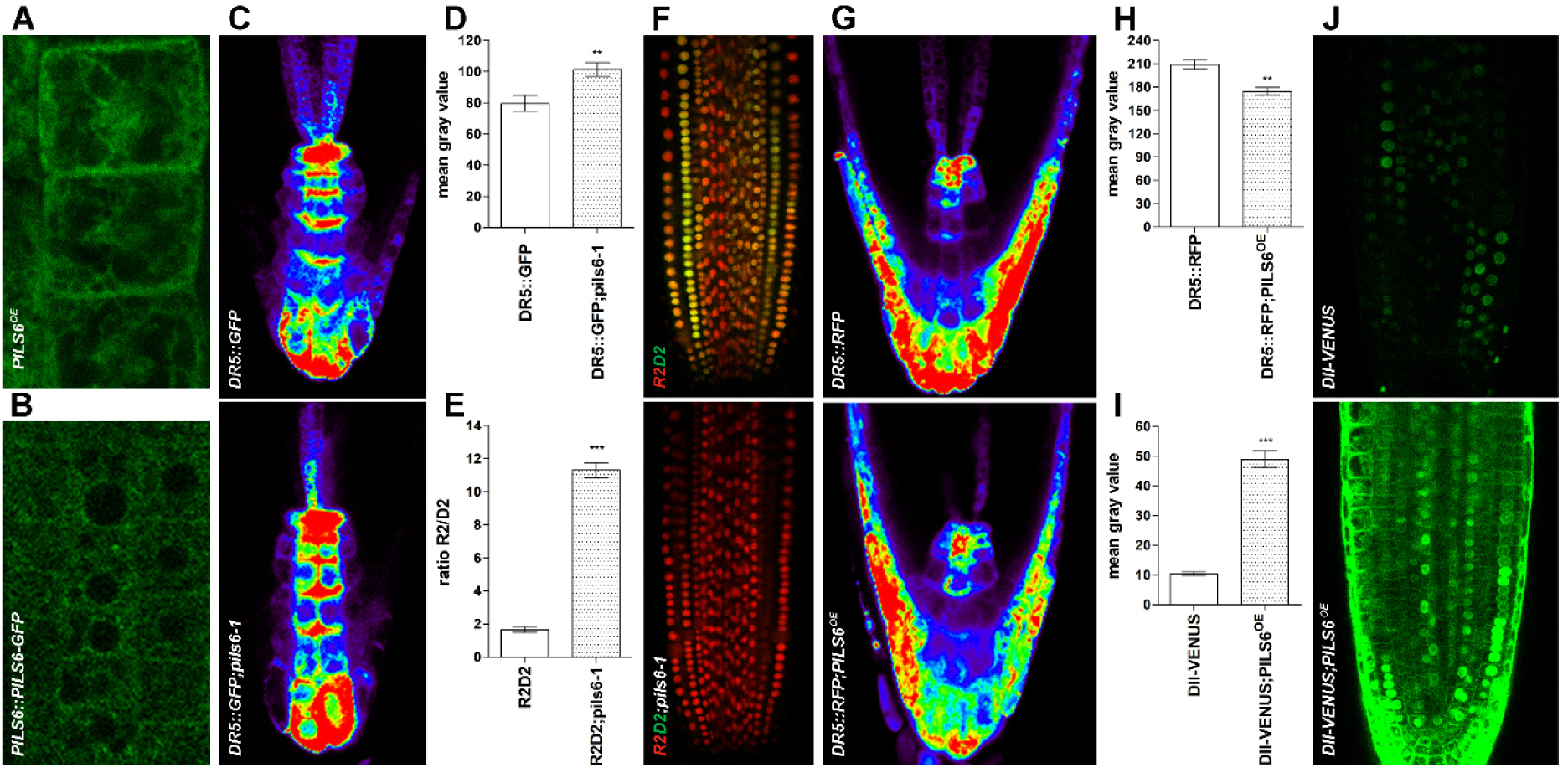
ER-localized PILS6 affects the nuclear availability of auxin. A-B PILS6 localizes to the ER. Confocal images of PILS6-GFP in the *PILS6*^*OE*^ (A) and *PILS6::PILS6-GFP* (B) show ER-like distribution in root meristem. C-J PILS6 affects auxin signalling. Pseudo-coloured confocal images (C, G) and fluorescence intensity quantification (D, H) of *pDR5rev::GFP* (C, D) or *pDR5rev::mRFP1er* (G, H) in the root meristems show stronger signal in the *pils6-1* (C, D) and weaker signal in the *PILS6*^*OE*^ (G, H) when compared to the wild type. Note that in the pseudo-coloured visualisation red is indicating high and blue low intensity. Merged confocal images of R2 (red) and D2 (green) expressing R2D2 marker (F) and quantification of R2/D2 ratio (E) show enhanced degradation of D2 signal in the *pils6-1* mutant. Simultaneous confocal imaging of DII-VENUS and PILS6-GFP (J) and quantification of DII-VENUS in the nuclei (I) show enhanced fluorescence in the *PILS6*^*OE*^ background (only the green signal in the nuclei of root cells) when compared to the wild type. n = 6 roots (D), 7-9 roots (E), 6 roots (H), and 50 nuclei from 5 roots per each genotype (I). *: p< 0.05, Student’s *t*-test.

To address if PILS6 also impacts on auxin biology, we initially assessed nuclear auxin signalling rates by using the auxin responsive promoter DR5 transcriptionally fused to GFP (*pDR5rev::GFP*; Benkova et al., 2003) or RFP (*pDR5rev::mRFP1er*; Marin et al., 2010). *pils6-1* mutant displayed higher *pDR5rev::GFP* expression in root apex (Figures 2C, 2D), indicating increased nuclear auxin signalling. In contrast, *pDR5rev::mRFP1er* was reduced in *PILS6*^*OE*^ expressing roots (Figures 2G, 2H), revealing decreased nuclear auxin signalling. Thus, similarly to PILS2, 3, and 5 proteins (Barbez et al., 2012; Beziat et al., 2017), PILS6 also negatively impacts on nuclear auxin signalling output.

PILS proteins have a presumed role in limiting auxin diffusion into the nucleus, due to an auxin compartmentalization mechanism at the ER (Barbez et al., 2012; Barbez et al., 2013; Barbez and Kleine-Vehn, 2013; Beziat et al., 2017). We therefore indirectly monitored PILS6-dependent nuclear availability of auxin, using the nuclear auxin input reporters DII-VENUS (Brunoud et al., 2012) and its related ratiometric version R2D2 (Liao et al., 2015). Nuclear DII/D2-domain displays auxin-sensitive degradation and, hence, the fluorescence of DII/D2-VENUS correlates inversely with the nuclear availability of auxin (Brunoud et al., 2012; Liao et al., 2015). *pils6-1* mutant displayed remarkably reduced D2 fluorescence (Figures 2E, 2F; Supplementary Figure 2E), indicating substantially higher nuclear auxin levels in the root apical meristem. On the other hand, *PILS6* overexpression specifically enhanced nuclear DII-VENUS signal intensities in the root apical meristem when compared to the control and auxin insensitive mDII-VENUS lines (Figures 2I, 2J; Supplementary Figure 2F), suggesting less nuclear availability of auxin. These findings suggest that the putative auxin carrier PILS6 limits the nuclear availability of auxin and subsequently affects auxin signalling and auxin-dependent root growth.

Notably, the impact of PILS6 on the nuclear availability of auxin appeared quantitatively much stronger than its impact on DR5-based auxin signalling rates. This observation may imply that additional processes modulate the nuclear auxin signalling output to compensate for the impact of PILS6 on nuclear auxin levels.

We recently showed that phyB-dependent light perception induces *PILS2* and *PILS3* expression and thereby impacts on auxin-dependent growth regulation in apical hooks (Beziat et al., 2017). Therefore, we assume that *PILS* genes could integrate other environmental cues into plant growth and accordingly looked for abiotic factors that may impact on *PILS6* expression. Publicly accessible transcriptomic data suggested that *PILS6* is particularly responsive to HT stress (Kilian et al., 2007; see also the eFP browser http://www.goo.gl/sG4BVw). Moreover, *pils6-1* mutants show an increase in auxin signalling (Figures 2C, 2D) and root growth (Figures 1D, 1E), which is reminiscent of wild type seedlings grown under HT conditions (Hanzawa et al., 2013; Wang et al., 2016; Fei et al., 2017; Martins et al., 2017). Thus, we speculated that PILS6 may also play a role in integrating environmental cues, such as temperature. To test this hypothesis, we initially analysed the response of the *PILS6* gene to HT by using a transcriptional fusion of PILS6 promoter sequence with the beta-glucoronidase (*GUS*) and *GFP* (*pPILS6::GUS-GFP*) reporter (Beziat et al., 2017). Notably, the same promoter sequence was used to complement the *pils6-1* mutant with PILS6-GFP (Supplementary Figure 2D). We readily detected *PILS6* expression in cotyledons, hypocotyls, and roots (Figure 3A; Supplementary Figures 3A, 3B), which is in agreement with endogenous transcript detection and cell-type specific, microarray-based predictions (Barbez et al., 2012; Feraru et al., 2012).

**Figure 3.**
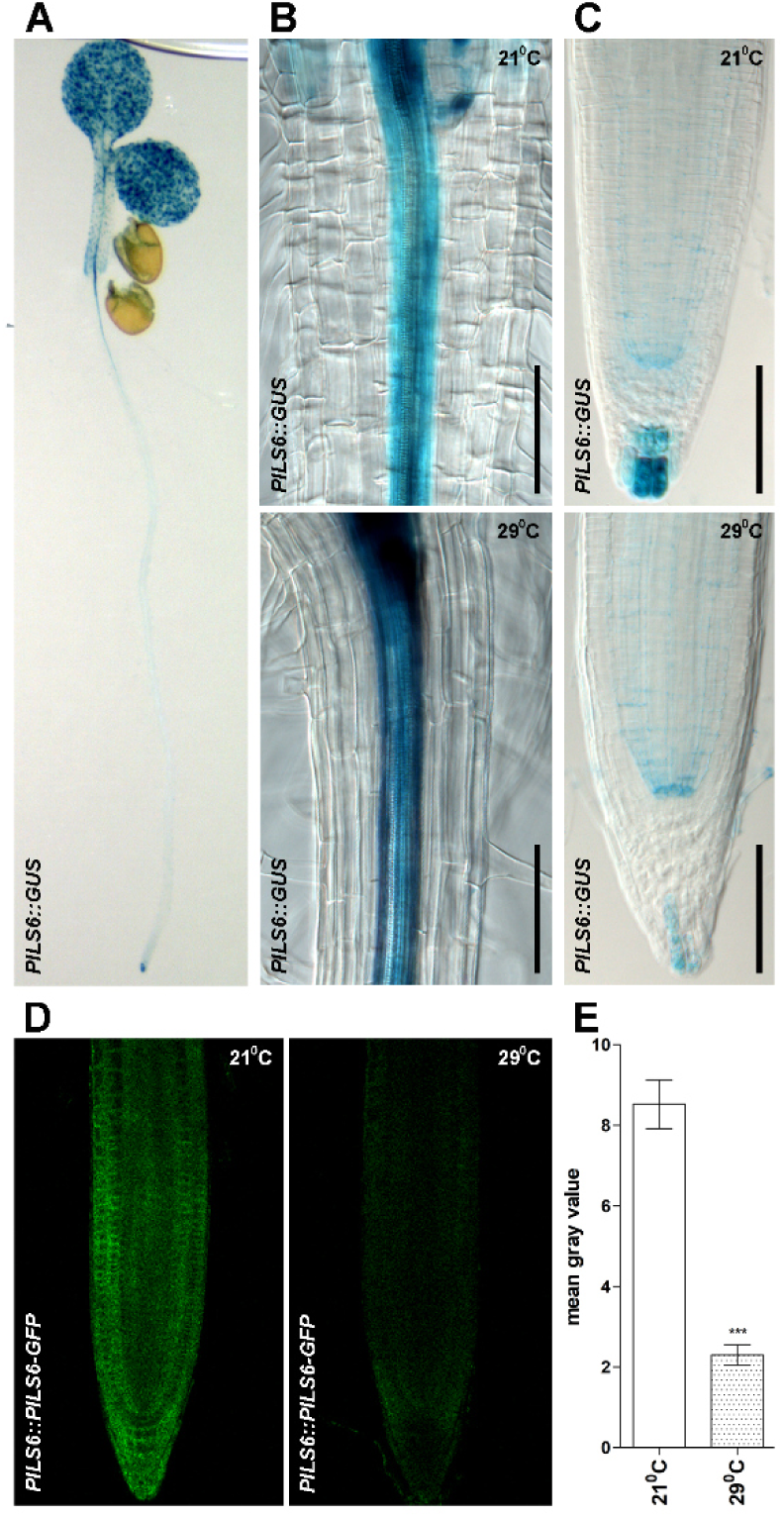
PILS6 expression and protein level are affected by HT. A Histochemical localisation of *pPILS6::GUS* shows gene expression in a 5-d-old seedling. B-E HT affects PILS6 expression (B, C) and protein level (D, E). Histochemical localisation of *pPILS6::GUS* activity shows that HT upregulates *PILS6* gene in the root region around the collet (B) and specifically downregulates the expression of *PILS6* in the root apex (C). Confocal images (D) and quantification (E) of PILS6-GFP fluorescence show that HT reduces the PILS6 protein levels in roots. Scale bars: 100 µm (B-C). n = 7 roots (E). *: p< 0.05, Student’s *t*-test.

HT treatment had an intriguing dual effect on *PILS6* gene expression. *PILS6* expression was prominently upregulated in some tissues (Figure 3B; Supplementary Figures 3C, 3D), but specifically downregulated in the root apex of seedlings grown under 29^o^C (Figure 3C). Importantly, HT treatment also diminished the PILS6 protein levels in the *PILS6::PILS6-GFP* root meristem (Figures 3D, 3E). Our data shows that HT reduces the transcription and protein level of PILS6, proposing a potential role of PILS6 in temperature-sensitive root growth.

To advance our understanding of HT-dependent regulation of PILS6, we imaged the constitutively-expressed PILS6-GFP in the root meristem of *PILS6*^*OE*^ seedlings grown under 21^o^C or 29^o^C. Despite its constitutive expression, PILS6-GFP fluorescence was much weaker when grown under HT (Figures 4A, 4B). Notably, the ER marker *35S::HDEL-GFP* line (Brandizzi et al., 2003) did not respond to HT (Supplementary Figures 4A, 4B), revealing a specific temperature-sensitive post-translational control of PILS6 protein levels.

**Figure 4.**
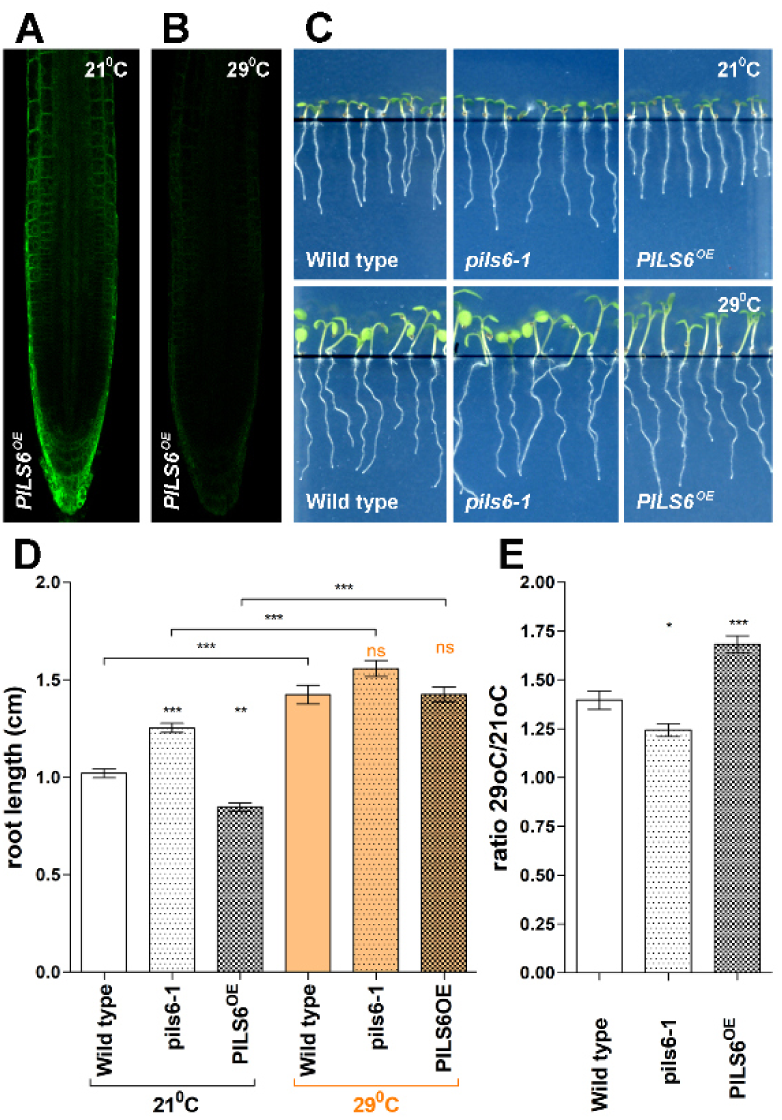
PILS6 regulates root growth under HT. A-B HT affects PILS6-GFP in the *PILS6*^*OE*^ roots. Confocal images show reduced PILS6-GFP fluorescence in the *PILS6*^*OE*^ roots grown under 29^o^C (B) when compared to 21^o^C (A). C-E PILS6 affects root growth under HT. Scanned images (C), measurements (D) and calculated ratio (E) of total root length show that *pils6-1* and *PILS6*^*OE*^ affect root growth under HT when compared to the wild type. n = 32-34 roots (D, E). ns: not significant; *: p< 0.05, one-way ANOVA.

These results show that HT does not only negatively regulate *PILS6* transcription, but in addition also the abundance of PILS6 protein in the root meristem. Given the importance of PILS6 for nuclear auxin input and consequently root organ growth, we assume that the HT-dependent regulation of PILS6 is developmentally important.

In agreement with its negative impact on PILS6, HT has been shown to increase auxin signalling rates (Hanzawa et al., 2013; Wang et al., 2016). Accordingly, we subsequently investigated if HT impacts upon PILS6-dependent root growth. As expected, wild type seedlings grown for five days under HT (29^o^C) showed increased primary root growth compared to control conditions (Figures 4C-4E). In agreement with its negative impact on PILS6 protein levels, HT abolished the short root phenotype of *PILS6*^*OE*^ (Figures 4C-4E). In relative comparison to wild type, *PILS6*^*OE*^ roots appeared more sensitive to the HT effect, showing a 20 % increase in the total root length in comparison to wild type roots (Figures 4C-4E). This finding suggests that HT-dependent repression of PILS6 protein abundance does impact upon root organ growth. Under standard growth conditions, the *pils6-1* mutant displayed enhanced root growth (Figures 1D, 1E, 4C-4E), which is actually partially phenocopied by wild type grown under HT (Hanzawa et al., 2013; Wang et al., 2016; Fei et al., 2017; Martins et al., 2017). In agreement, *pils6-1* mutant roots showed a relative reduction in HT-induced root growth when compared to wild type seedlings (Figures 4C-4E). Our set of data indicates that HT modulates PILS6 abundance, which consequently impacts on root organ growth.

Our data shows that, similar to PILS2, 3 and 5, PILS6 also functions at the ER to repress auxin signalling (Barbez et al., 2012; Barbez et al., 2013; Beziat et al., 2017). Intriguingly, PILS2, 3, 5 and 6 belong to distinct subfamilies (Feraru et al., 2012), which suggests that all PILS subfamilies have a conserved function in restricting auxin signalling in *Arabidopsis*. Importantly, here we show that PILS6 controls the nuclear availability of auxin, which is in agreement with our model, assuming that PILS proteins transport auxin into the ER lumen to prevent its diffusion into the nucleus. We recently showed that light perception triggers the expression of *PILS2* and *3*, allowing this environmental cue to induce an auxin signalling minimum during apical hook development (Beziat et al., 2017). In contrast, here we show that an increase in ambient temperature negatively impacts on PILS6 abundance, which consequently modulates root organ growth. Accordingly, we assume that PILS proteins have general importance for integrating environmental cues into growth programs.

PIF4-dependent auxin biosynthesis integrates elevated temperature with growth in shoots, but not in roots (Martins et al., 2017). Accordingly, the role of auxin in elevated temperature-induced root growth remained controversial. We illustrate that HT strongly diminishes the abundance of PILS6, which notably correlates with an increase in auxin signalling (Hanzawa et al., 2013; Wang et al., 2016). It remains to be seen how temperature is sensed and PILS6 abundance is decreased. Nevertheless, we propose that PILS6 is part of a novel mechanism, linking HT to auxin responses in roots.

## Material and methods

### Plant material

*Arabidopis thaliana* ecotype Columbia 0 was used as the wild type. *35S::PILS6-GFP* (*PILS6*^*OE*^; Barbez et al., 2012)*, PILS6::GUS-GFP* (Beziat et al., 2017), *pDR5rev::GFP* (Benkova et al., 2003), *pDR5rev::mRFP1er* (Marin et al., 2010), R2D2 (Liao et al., 2015), DII-VENUS (Brunoud et al., 2012), mDII-VENUS (Brunoud et al., 2012), and *35S::HDEL-GFP* (Brandizzi et al., 2003) have been described previously. *pils6-1* (SALK_130335.46.50) and *pils6-2* (SALK_074172.15.55) were identified in the publically available T-DNA collections (Alonso et al., 2003) and ordered from NASC.

### Growth conditions

Seeds were sterilized either i) overnight with chlorine gas, followed by few hours aeration or ii) 1-2 min with 70% ethanol, followed by drying. Seeds were afterwards plated on one single line in the upper part of Petri dishes containing 50 ml solidified agar medium, made of 0.8% agar, 0.5x Murashige and Skoog (MS) medium, and 1% sucrose (pH 5.9). Prior to germination, the seeds were stratified for 2-3 days in the dark at 4^o^C. Then, seedlings were grown on vertically-oriented plates under standard growth conditions (plant cabinet equipped with above placed cool-white fluorescent bulbs and set at about 140 µmol/m^−2^s^−1^, long day photoperiod, and 21^o^C).

For HT-related experiments, two growth cabinets were equipped with overhead LED cultivation lights (Ikea, 703.231.10), at an irradiance of 170 µmol/m^−2^s^−1^, long day photoperiod, and set at 21^o^C (control) or 29^o^C (HT treatment). After stratification, the seeds were allowed to germinate for 24 hours at 21^o^C, and then grown on vertically-oriented plates for 4-5 days under 21^o^C (control) or 29^o^C (HT).

### Quantification of cotyledons, rosettes and root phenotype

For cotyledons and rosette measurements, seedlings were grown initially on vertically-oriented plates for 5 days, then turned to a horizontal position, and allowed to grow for 3 (cotyledons) or 7 (rosettes) days more. For root measurements, seedlings were grown for 5-7 days on vertically-oriented plates. Plates were scanned with Epson Perfection V700 scanner. Rosette area and root length were measured by using ImageJ 1.41 software (http://rsb.info.nih.gov/ij/).

### Quantification of PC and stomatal size

8-day-old seedlings grown on vertically-oriented plates as described above were harvested in the morning, usually 2-3 hours after lights turned on in the cabinet. Cotyledons were harvested directly into 95% ethanol and fixed for 3-4 hours at room temperature. Then samples were incubated successively in ethanol dilutions of 70%, 50% and 20% (each step 10 minutes) and cleared overnight in chloral hydrate (1:3:8; glycerol:water:chloral hydrate). Cleared cotyledons were placed on slides with abaxial face upward and imaged under a Zeiss microscope, with a 20x objective. Images were always taken in the same region of the cotyledon and used in ImageJ for quantification of PC area and stomatal size index. The stomatal size index represents the length values multiplied by width values.

### GUS staining

This was performed as described previously (Beziat et al., 2017b). Prior to staining, 5-day-old seedlings were fixed in 90% cold acetone for 1 hour on ice, then washed in phosphate buffer for 1 hour at room temperature. Seedlings were stained overnight.

### Cloning

This was performed as described in Barbez et al. (2012). A promoter fragment of 1.2 kb was amplified with primers listed in Supplementary Table 1 and subsequently cloned by Gateway recombination, firstly into pDONR221, and then into pKGWFS7.0 (Karimi et al., 2002). Transformed lines were selected on Kanamycin. To generate *PILS6::PILS6-GFP*, the full genomic fragment was cloned into the pDONR221 and the promoter region into the pDONR-P4P1, by using the primers listed in Supplementary Tabel 1. These entry clones and the GFP-containing entry clone were subsequently transferred to the gateway-compatible destination vector pK7m34GW,0 (Karimi et al., 2002). Transformed lines were selected on Kanamycin.

### Confocal imaging

Imaging was performed on 5-day-old seedlings, by using a Leica TCS SP5 confocal microscope, equipped with HyD in addition to the standard PMTs. The signal intensity (mean gray value) of the marker lines was quantified by using “Quantify” tool of SP5 software (LAS AF Lite). DII-VENUS signal in nuclei was quantified by defining a round Region Of Interest (ROI) in the middle of the nucleus, while for all the other markers we defined rectangle ROIs in the most representative region of the root.

### Data analysis

We used Excel to organize data and GraphPad Prism 5 software for statistical analysis and graphing. For statistical analysis of our raw data, we used one-way ANOVA and Tukey’s multiple comparison test for the experiments with several genotypes (e.g. wild type control, *pils6* mutants and *PILS6*^*OE*^), and Student’s *t*-test for the experiments with two genotypes/conditions (e.g fluorescence intensity in two different backgrounds or conditions).

Representative experiments are shown.

## Acknowledgements

We are grateful to J. Friml, A. Maizel, T. Vernoux, and D. Weijers, for providing the published marker lines; the BOKU-VIBT Imaging Center for access and M. Debreczeny for expertise; L. Mach and R. Strasser for sharing equipment. This work was supported by the Vienna Science and Technology Fund (WWTF) (to J.K.-V.), Austrian Science Fund (FWF) (Projects: P26568-B16 and P26591-B16 to J.K.-V.), European Research Council (AuxinER - ERC starting grant to J.K.-V.), European Molecular Biology Organization (EMBO) (ALTF 795-2012 to E.F.) and FWF-Hertha Firnberg (T728-B16 to E.F.).

